# iPheGWAS: an intelligent computational framework to integrate and visualise genome-phenome wide association results

**DOI:** 10.1101/2022.03.05.483121

**Authors:** Gittu George, Yu Huang, Sushrima Gan, Aditya S. Nar, Jason Ha, Radha Venkatesan, Viswanathan Mohan, Huan Wang, Andrew Brown, Colin N. A. Palmer, Alex S. F. Doney

## Abstract

Estimating the genetic correlations by LDSC is computationally demanding and visualising multiple GWAS results along with their genetic relationships is restricted. This study developed iPheGWAS, a novel approach which applied hierarchical clustering to GWAS summary statistics to (i) calculate their genetic relatedness, and (ii) enable three-dimensional visualisation of multiple ordered GWAS plots. Simulation and real-world data analysis demonstrated that when investigating genetic relationships among multiple phenotypes, iPheGWAS can deliver comparable results with LDSC but with 8 times faster computational speed. It can also provide novel findings in studying genetically-correlated comorbidities, such as mental illness and rheumatoid arthritis.

## Background

The development from genome-wide association studies (GWAS) to phenome-wide association study (Phewas), which test genetic variants against large numbers of phenotypes, has given us the potential to explore in depth the complex relationships between genomes and phenomes. The incorporation of Electronic Health Records systems (EHRs) has garnered widespread acceptance across a plethora of genomic medicine applications. In our previous work, we have developed ‘PheWAS’, a software package for visualising plots of genome-wide data, e.g. Manhattan plots, across a number of phenotypes as a three-dimensional landscape (1). This makes it a great tool for identifying SNPs with pleiotropic effect and exploring local genetic correlations and colocalization (1). However, one limitation of ‘PheWAS’ is that carrying out statistical analysis on its required data form is difficult, which limits its use in identifying the overall genome-phenome relationships

One important relationship between pairs of traits is their genetic relatedness, known as ‘genetic correlation’. Genetic correlation between traits can be estimated when multiple GWAS results are available. As a quantitative population parameter it provides a measure of the similarity of the genetic architecture of the two traits (2,3). Estimates of genetic correlation can show evidence of pleiotropic action of genetic variants, informing on possible shared causal processes (4–6).

Several methods have been developed for estimating genetic correlations - some of these methods require individual level data, such as Genome-wide Complex Trait Analysis (GCTA) (7), MTG2 (8) and BOLT-REML (9), which based on Genome-based Restricted Maximum Likelihood (GREML) method. Recently, there is enormous demand of analysing genetic correlation with GWAS summary statistics due to the ease of use, computational speed and feasibility even when individual level data are not available (10–13). Furthermore, concerns regarding the privacy of individual level data are mitigated with the use of summary statistics for estimating genetic correlation rather than traditional approaches (14). Among these methods, the LD Score Regression (LDSC) (13) is the most widely used. However, none of aforementioned methods are able to estimate and present the genetic relationship among several traits in one single test or plot.

Hierarchical clustering is a machine learning method that assembles traits by their similarity. Previous application of hierarchical clustering in genomic data is limited. However, in omic studies, such as transcriptomic (15) and proteomics studies (16), hierarchical clustering is widely used. We propose that, by selecting proper genetic features from the GWAS summary statistics, hierarchical clustering could efficiently generate cluster of traits that are genetically correlated.

We previously developed a tool (PheGWAS) to visualise ‘many variants-many phenotypes’ in a single plot. However, PheGWAS is unable to reflect the genetic correlation information. In this paper, we have developed a heuristic approach based on hierarchical clustering, in order to **1) develop a method that can efficiently cluster or organize multiple traits according to their genetic similarity in one single test; 2) develop a tool to assist the visualisation of genome-wide data plots across a number of phenotypes with integrative genetic relationship information.** We compared our method with the gold-standard technique LDSC and applied the heuristic method to the PheGWAS landscape and re-introduced the landscape as an intelligent PheGWAS landscape (iPheGWAS).

## Results

### Results for simulation data

In the simulation studies with 500 causal SNPs, it was observed that when the genetic correlations between the pair of traits and the heritability for each trait were both high as in Arc 1 and Arc 4, and all of the settings of iPheGWAS performed well (Table 2). The overall success rate dropped when the heritability was low (Arc 5 in Table 2). In scenario when the genetic correlations were as low as 0.1, though heritabilities were high, the overall success rate was low (Acr 2 and Acr 3 in Table 2). iPheGWAS outperformed LDSC when heritability or genetic correlation was low (Acr 5, Acr 2, Acr 3 in Table 2), while within iPheGWAS, the ‘pairwised’ set up had lowest success rate. (Table 2)

**Table 1.** Five different genetic architecture setups for simulation set 1 and 2.

|  | Genetic correlations (rg) |  |  | Heritability (h<sup>2</sup>) |  |  |  |
| --- | --- | --- | --- | --- | --- | --- | --- |
|  | a-b | c-d |  | a | b | c | d |
| Arc 1 | 0.3 | 0.7 |  | 0.5 | 0.7 | 0.5 | 0.7 |
| Arc 2 | 0.1 | 0.7 |  | 0.5 | 0.7 | 0.5 | 0.7 |
| Arc 3 | 0.1 | 0.2 |  | 0.5 | 0.7 | 0.5 | 0.7 |
| Arc 4 | 0.7 | 0.8 |  | 0.5 | 0.7 | 0.5 | 0.7 |
| Arc 5 | 0.3 | 0.7 |  | 0.1 | 0.2 | 0.1 | 0.2 |

rg: genetic correlation between pairs of traits; h<sup>2</sup>: heritability.

**Table 2.** Success rate of the simulation study with 500 causal SNPs.

|  | Arc 1 | Arc 2 | Arc 3 | Arc 4 | Arc 5 |
| --- | --- | --- | --- | --- | --- |
| Non-pairwise(spearman) | 10/10 | 8/10 | 8/10 | 10/10 | 9/10 |
| Non-pairwise(pearson) | 10/10 | 8/10 | 8/10 | 10/10 | 9/10 |
| Pairwise (spearman) | 10/10 | 7/10 | 6/10 | 10/10 | 9/10 |
| Pairwise (pearson) | 10/10 | 7/10 | 5/10 | 10/10 | 10/10 |
| Skelton SNP's (spearman) | 10/10 | 7/10 | 8/10 | 10/10 | 10/10 |
| Skelton SNP's (pearson) | 10/10 | 7/10 | 8/10 | 10/10 | 10/10 |
| LDSC | 9/10 | 4/10 | 4/10 | 10/10 | 6/10 |

In the simulation studies with 60 causal SNPs, the overall success rate was lower compared with simulation study with more causal SNPs. The overall trend was similar as the previous simulation study, that is with the decrease of genetic correlation and heritability, the overall success rate declined. It was difficult for LDSC to distinguish pairs of traits when the inter genetic correlation was as low as 0.1 (Arc 2 and Arc 3 in Table 3) or when the heritability was as low as 0.1 (Arc 5 in Table 3). Within iPheGWAS, the ‘Skelton SNP’ set up had relatively higher success rate.(Table 3)

**Table 3.** Success rate of the simulation study with 60 causal SNPs. Each row represents the feature and correlation method used and each of the columns represents the various genetic architectures.

|  | Arc 1 | Arc 2 | Arc 3 | Arc 4 | Arc 5 |
| --- | --- | --- | --- | --- | --- |
| Non-pairwise(spearman) | 9/10 | 5/10 | 4/10 | 10/10 | 9/10 |
| Non-pairwise(pearson) | 9/10 | 5/10 | 4/10 | 10/10 | 9/10 |
| Pairwise (spearman) | 9/10 | 5/10 | 5/10 | 10/10 | 8/10 |
| Pairwise (pearson) | 10/10 | 5/10 | 5/10 | 10/10 | 6/10 |
| Skelton SNP's (spearman) | 9/10 | 6/10 | 5/10 | 10/10 | 9/10 |
| Skelton SNP's (pearson) | 9/10 | 5/10 | 5/10 | 10/10 | 9/10 |
| LDSC | 8/10 | 5/10 | 1/10 | 10/10 | 3/10 |

### iPheGWAS set up finalization

From the simulation study, it was found that the error rate of ‘Pairwise’ method was higher, hence ‘Non-pairwise’ and ‘Skeleton SNP’ methods were considered. Since using skeleton SNPs as features were based on pre-defined set of SNPs, the number of features for each trait was fixed regardless the number of traits being analysed. In the ‘Non-pairwise’ method, the number of features would increase in proportion to the number of traits being analysed, which is not computationally efficient. Hence, the ‘Skeleton SNP’ method was selected. Even though Pearson and Spearman correlation methods performed equally, Pearson’s correlation method was preferred, as Spearman correlation would discard information contained in highly significant genetic signals.

Taking into consideration the validity in clustering, we concluded that using skeleton SNPs as features, Average method as the linkage criteria and Pearson correlation method for generating the distance matrix provided with lowest error rates.

### Real world data validation: Study 1-Traits selected from UKBB (iPheGWAS)

The dendrograms for study 1 using our method and LDSC results were represented in Figure 2. In iPheGWAS it was observed that for the anthropomorphic domain, ‘trunk’ and ‘mass’ have the shortest height indicating they were the closest and were clustered first (clade 1); for the mental health domain, ‘neuroticism score’ and ‘mood swing’ clustered first (clade 2); for food related traits, the leaves ‘food weight’ and ‘potassium’ clustered first (clade 3), and for the cardiology domain, the intra correlations among traits were less strong compared to other domains, hence they clustered later- ‘cholesterol lowering’ and ‘no heart problem’ formed the first cluster (clade 5) and ‘pain throat’ and ‘atherosclerosis’ formed the second cluster (clade 6). We found that the relationship between ‘portion size’ and anthropomorphic traits was closer compared to its original domain (Figure 2 a). More details describing the dendrogram were included in supplementary results 1.

**Figure 1.**
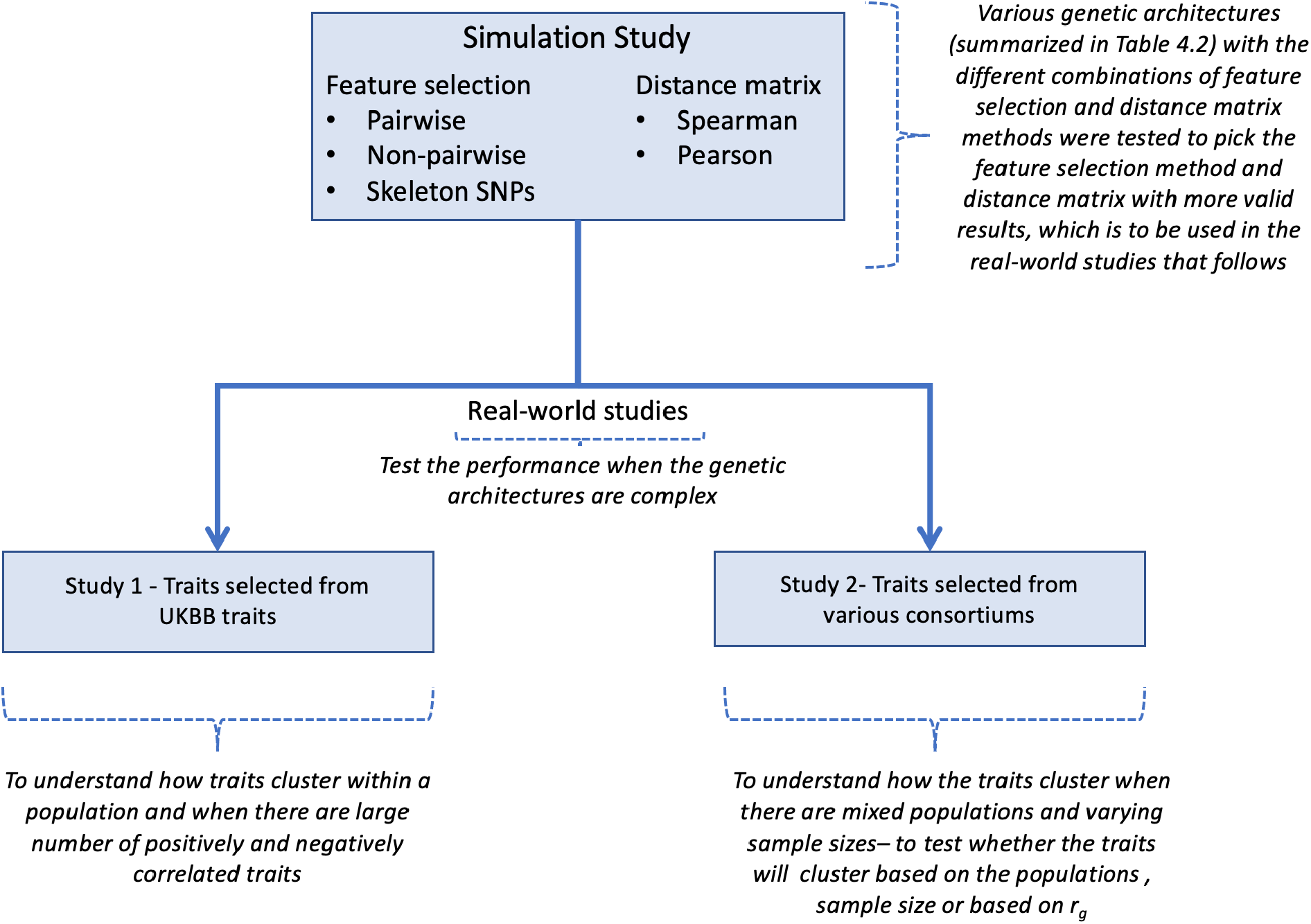
Flowchart explaining the steps involved in our study.

**Figure 2:**
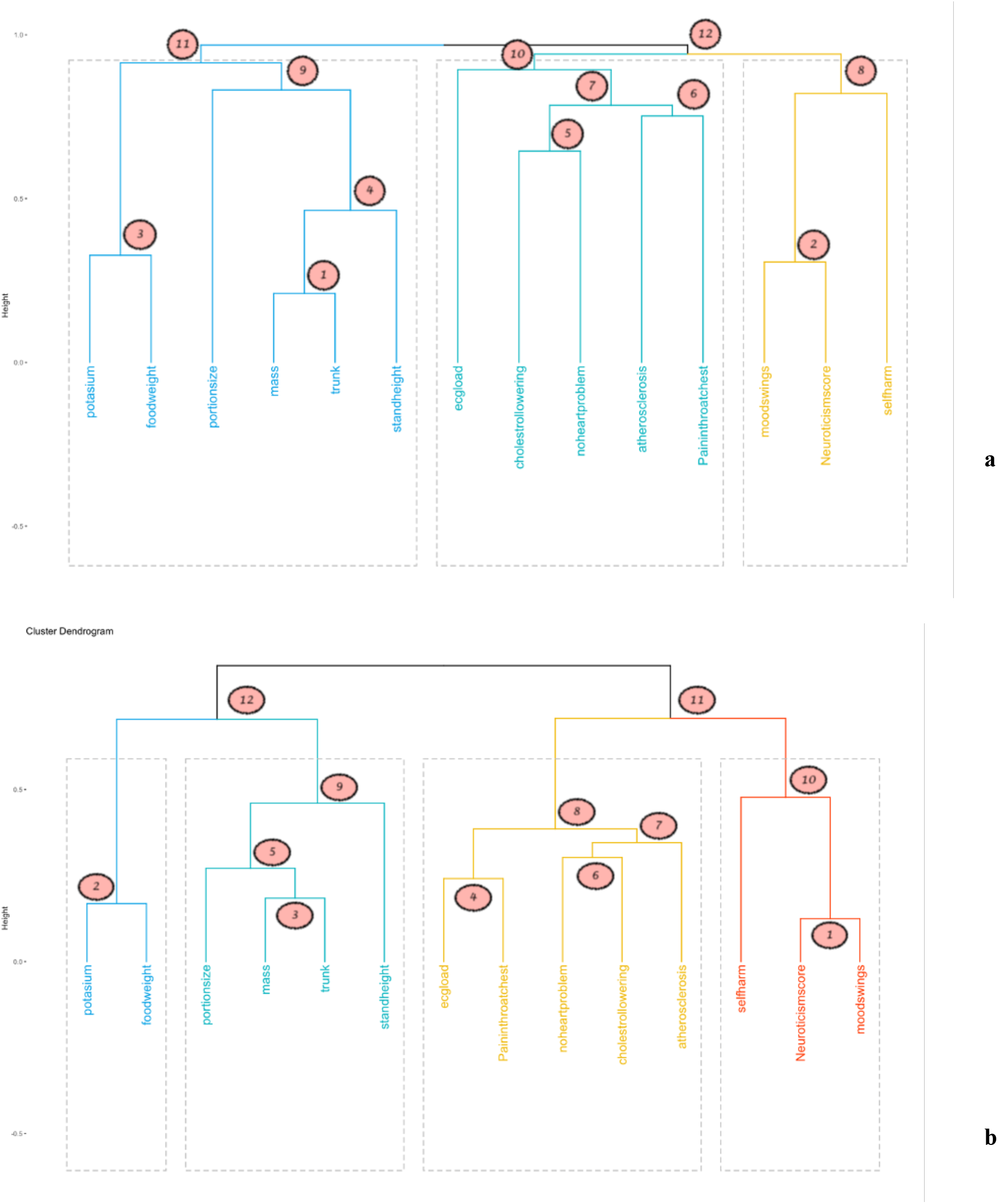
Dendrogram representing the hierarchical clustering for study. 1. a) Dendrogram generated by iPheGWAS. b) Dendrogram generated using LDSC matrix. Grey dotted box represent different clusters.

The tree based internal validation was high (cophenetic correlation coefficient = 0.948), suggesting the dendrogram represented the structure of the data well. The optimal number of clusters obtained from the Silhouette method suggested 3 clusters (Supplementary figure 3). Based on these results three clusters were suggested by iPheGWAS, and the elements included in each cluster were shown in Figure 2 a,.

### Real world data validation: Study 1- Traits selected from UKBB (LDSC)

In dendrogram generated using LDSC matrix, for anthropomorphic, mental health and food related domain respectively, the clusters in the first level were the same although the intra cluster relatedness was slightly different; for the cardiology domain, ‘ECG load’ and ‘pain in throat/chest’ were found to be more similar unlike in iPheGWAS and these formed a cluster (clade 4), leaving ‘atherosclerosis’ joining the cluster as the second level. In common with iPheGWAS, ‘portion size’ and anthropomorphic traits were closer, even closer than ‘standing height’ (Figure 2 b). More details describing the dendrogram are included in supplementary results 2.

The optimal number of clusters obtained from Silhouette method suggested 4 clusters (Supplementary figure 4). Based on these results four clusters were suggested by LDSC matrix, and the elements included in each cluster are shown in Figure 2 b.

### Real world data validation: Study 1- Comparison of the dendrograms and the clusters

A Tanglegram plot is shown in Figure 3. The entanglement coefficient between the two dendrograms was very low (0.09), suggesting the alignment of both the two trees was good (Figure 3).The tree based external validation cophenetic correlation coefficient and baker’s gamma coefficient were high (0.85 and 0.99 respectively).

**Figure 3:**
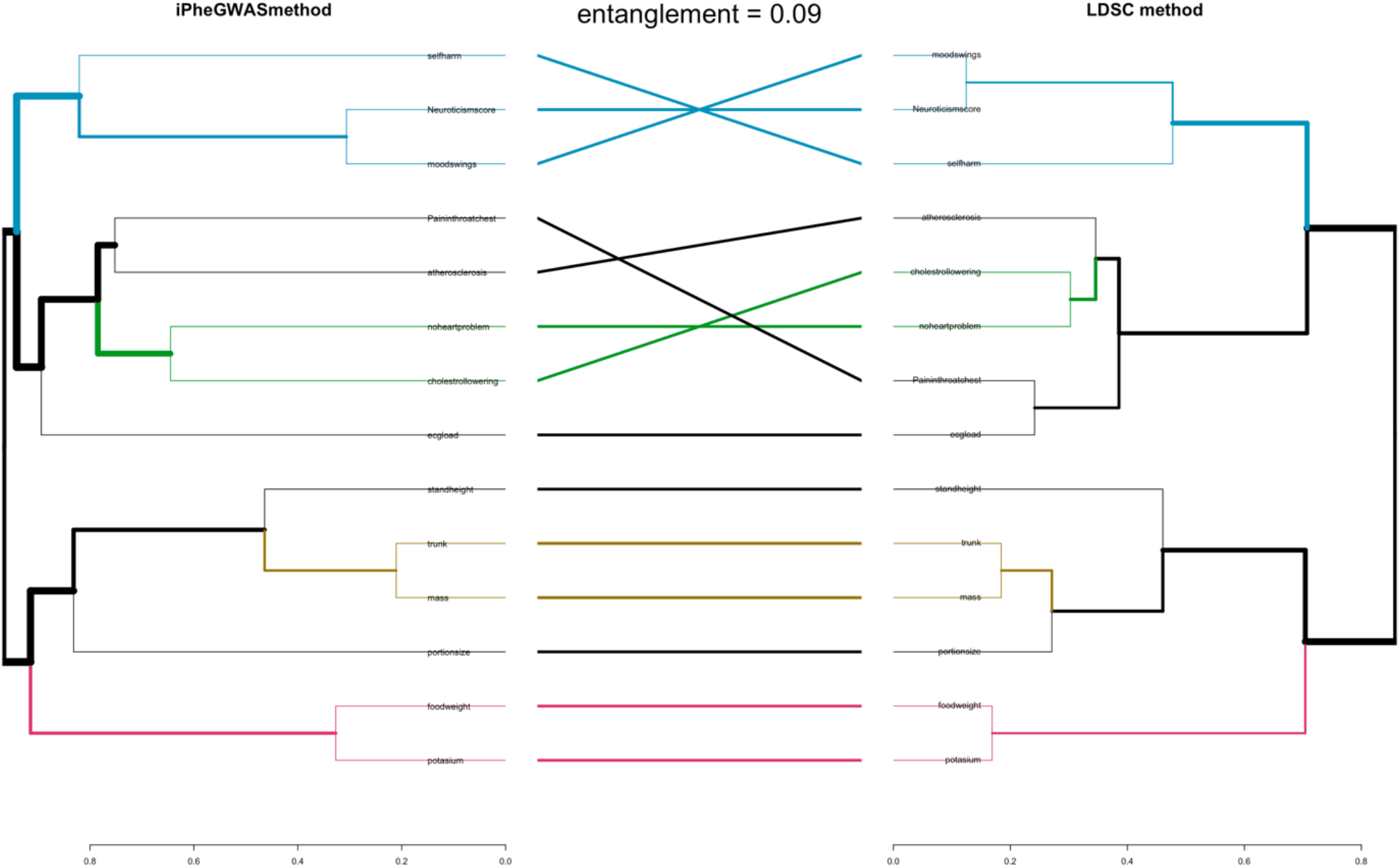
Tanglegram comparing the dendrograms obtained from iPheGWAS and LDSC methods for study 1.

Although the optimal number of clusters suggested for iPheGWAS was three, four clusters were suggested for the LDSC method. The Rand Index was 0.91 and the Adjusted Rand Index was 1, suggesting the similarity between the clusters is very high.

### Real world data validation: Study 2-Traits selected from various consortiums (iPheGWAS)

The dendrograms for study 2 using iPheGWAS and LDSC results were represented in Figure 4. In iPheGWAS it was observed that the five traits from the psychiatry consortiums were clustered indistinguishably as clade 1. Interestingly, ‘Rheumatoid Arthritis’ from autoimmune diseases joined psychiatric traits as the second level (clade 2), suggesting a closer relationship between them; among the rest autoimmune diseases, ‘Ulcerative Colitis’ and ‘IBD’ clustered first; for anthropomorphic traits, ‘waist hip ratio’ and ‘BMI’ clustered first (clade 5), followed by a second cluster formed by ‘birthweight’ and ‘height cluster’ (clade 6). Another finding is, ‘education’ was found to be more similar to ‘waist hip ratio’ and ‘BMI’ compared to ‘height’ and ‘birth weight’ (Figure 4 a). More details describing the dendrogram were included in supplementary results 3.

**Figure 4:**
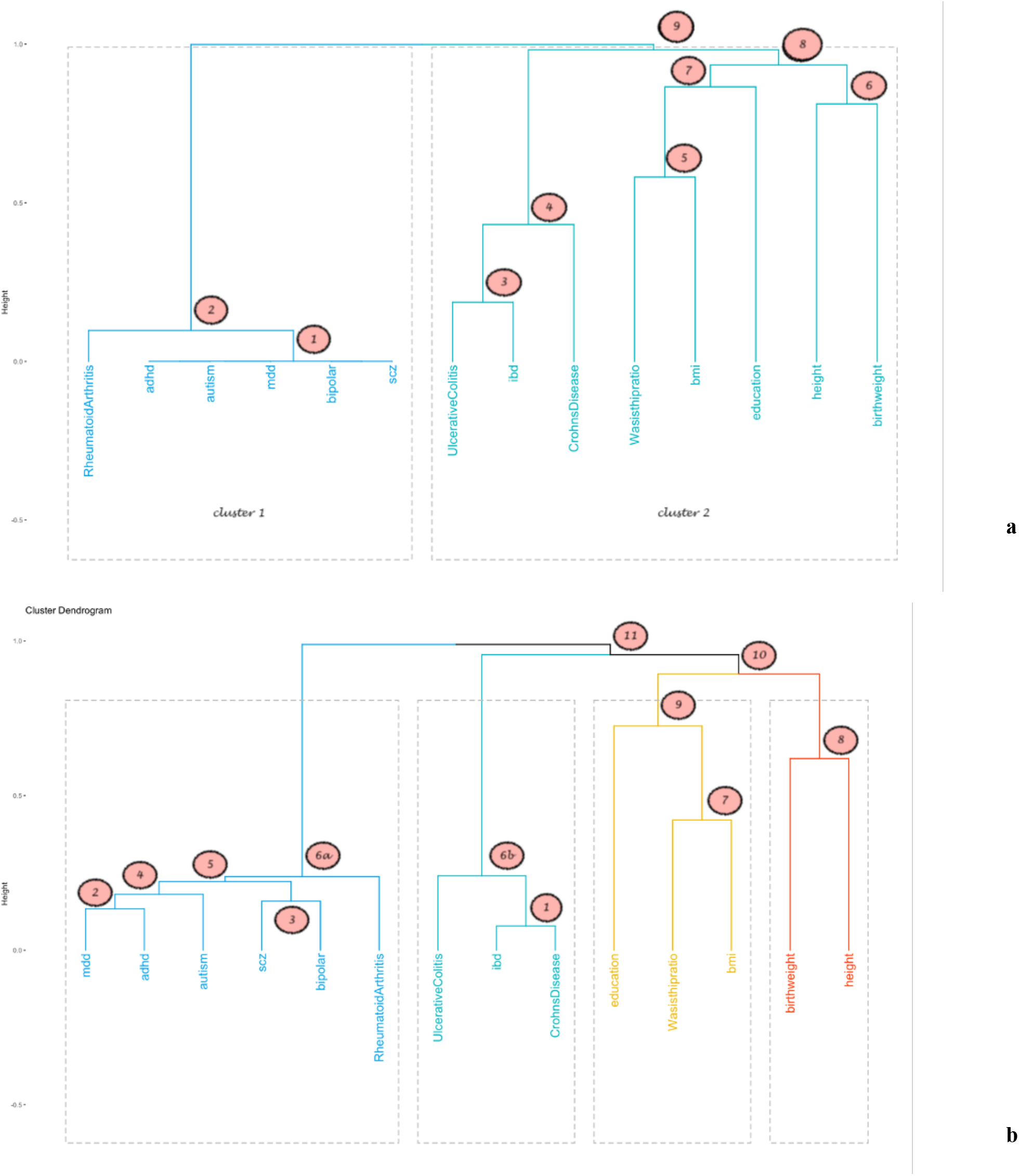
Dendrogram representing the hierarchical clustering for study 2. a) Dendrogram generated by iPheGWAS. b) Dendrogram generated using LDSC matrix. Grey dotted box represent different clusters.

The tree based internal validation was very high (cophenetic correlation coefficient=0.995). The optimal number of clusters obtained from Silhouette method suggested two clusters (Supplementary figure 5). The two clusters identified by iPheGWAS were - cluster 1 containing the psychiatry traits and rheumatoid arthritis and cluster 2 containing all the other traits (Figure 4 a).

### Real world data validation: Study 2-Traits selected from various consortiums (LDSC)

The dendrogram for study 2 using LDSC is represented in Figure 4 b. It is observed that ‘IBD’ and ‘Crohns Disease’ from autoimmune diseases consortium clustered first, unlike the psychiatry traits in iPheGWAS (clade 1). The five psychiatry traits were clustered closely by pairs and ‘rheumatoid arthritis were also sub-branches of the same clade, similar to iPheGWAS (clade 2 to 6a). Although ‘BMI’ and ‘Waist hip ratio’ clustered together similar to iPheGWAS and ‘education’ joined this cluster as a simplicifolius- ‘education’ was found to be closer to ‘Waist hip ratio’ (clade 9). ‘Birth weight’ and ‘height’ clustered together like in iPheGWAS but ‘birth weight’ was found to be closer to ‘BMI’.

The optimal number of clusters obtained from Silhouette method suggested 4 clusters (Supplementary figure 6). Based on these results four clusters were suggested by LDSC matrix, and the elements included in each cluster were shown in Figure 4 b.

### Real world data validation: Study 2-Comparison of the dendrograms and the clusters

Tanglegram plot was shown in Figure 5. The entanglement coefficient between the two dendrograms was very low (0.09), suggesting the alignment of the two trees was good (Figure 5). The tree based external validation cophenetic correlation coefficient and baker’s gamma coefficient were 0.97 and 0.997 respectively.

**Figure 5:**
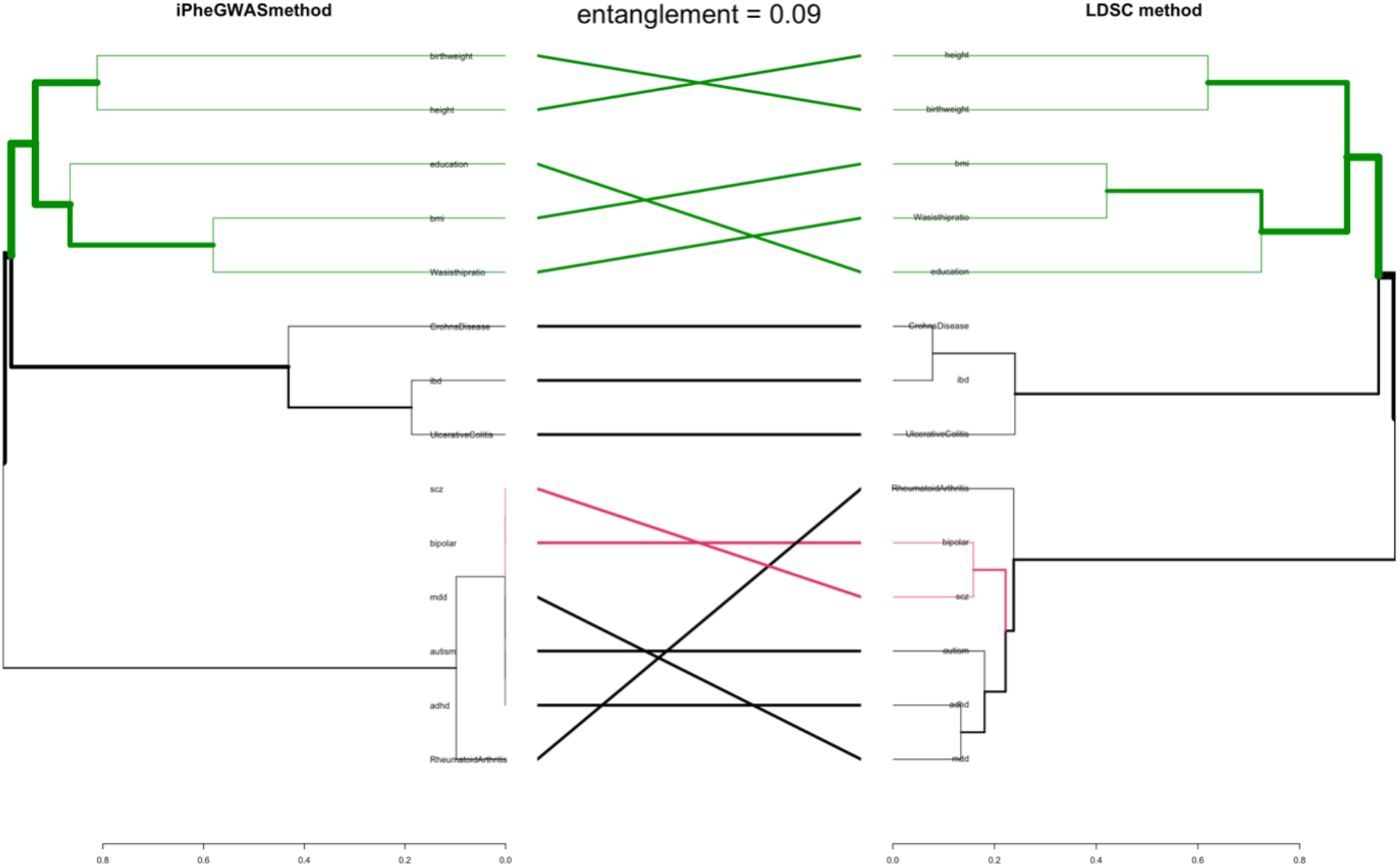
Tanglegram comparing the dendrograms obtained from iPheGWAS and LDSC methods for study 2.

Although the optimal number of clusters suggested for iPheGWAS was two, four clusters were suggested for the LDSC method. The Rand Index was 0.77 and the Adjusted Rand Index was 0.53.

### Demonstration of iPheGWAS on Selected Traits from Study 2

Six traits from autoimmune diseases and anthropomorphic traits were selected for demonstration. Figure 6 showed the PheGWAS landscape before and after applying the heuristic approach. After applying iPheGWAS, multiple traits were ordered and clustered according to their genetic relatedness, and the ‘peaks’ on the iPheGWAS plot were linked.

**Figure 6:**
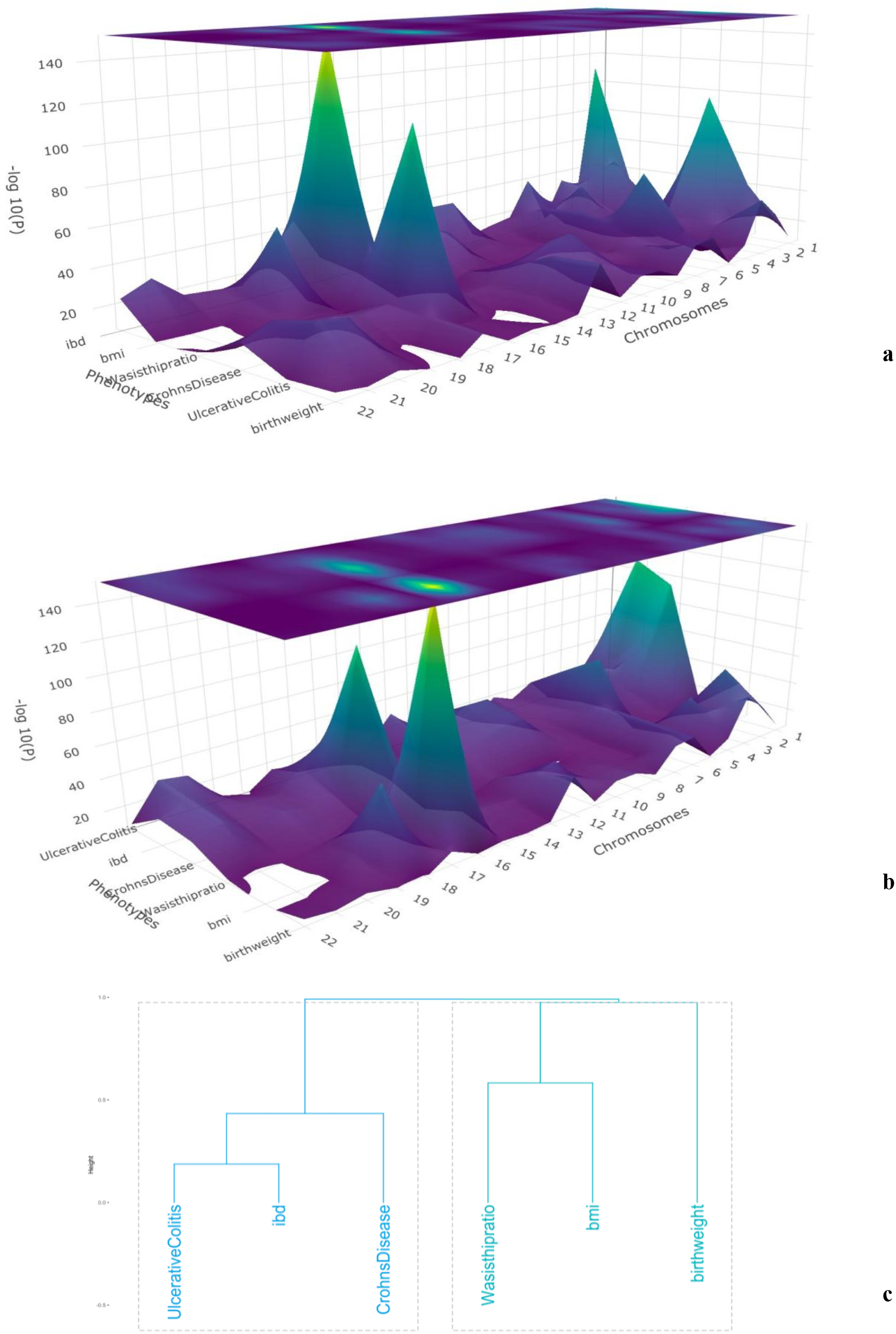
Demonstrating of iPheGWAS plot. The chromosome numbers were in X axis, Y-axis is different traits were aligned in Y axis and -log10 GWAS P values were in Z axis. a) PheGWAS plot before ordering; b) PheGWAS plot after ordering (iPheGWAS plot); c) dendrogram illustrating the relationship among traits.

### Computation efficiency

Under High Performance Computing (HPC) cluster working environment, iPheGWAS took 1.5 minutes for computation as compared to 12 minutes in LDSC for ordering 14 traits based on their genetic correlations.

## Discussion

In this study, we have introduced a novel heuristic approach to incorporate ordering of multiple GWAS traits based on their genetic similarity in the three-dimensional landscape. To the best of our knowledge, this is 1) the first visualisation tool for displaying and analyzing genetic similarity for multiple GWAS traits; and 2) the first approach that simultaneously computes genetic similarity for multiple traits using GWAS summary statistics compared to the traditional pairwise approach of computation. Our method suggests that: 1) the order of clustering of traits computed were similar with the order produced by the genetic correlations calculated by the LDSC method for most of the traits; 2) this method not only offers ease of use but is computationally efficient.

Our method provides several insights about the pattern of genetic relationship among phenotypes, both local and overall genetic correlations. Since iPheGWAS provides an ordered or clustered visualisation of multiple traits that are genetically similar, an easy visual appreciation of the overall genome-wide landscape provides initial clues about shared genetic effects across multiple phenotypes. By aligning two related traits side by side, clues about pleiotropy can be easily obtained for further investigation. Furthermore, iPheGWAS also assists the process of selecting traits for a multi-trait analysis genome-wide association studies (MTAG) (24) to improve power for detecting genetic variants contributing to disease risk.

A recent study applied hierarchical clustering to gene level association statistics to detect shared genetic architecture among multiple traits(25). Instead, we used full GWAS summary statistics as input for iPheGWAS. There are several advantages of using full GWAS summary statistics. Firstly, the number of features selected by iPheGWAS were greater, they covered gene coding region as well as non-coding regions where eQTLs might lie, and this might increase the power for detecting genetic correlation. As shown, in the simulation studies, iPheGWAS performed with less error rate when ‘skeleton SNPs’ were used as features. In general complex traits are more polygenic in nature (26), thus a probable reason for this could be because greater number of features are needed to capture the genetic structure of complex traits. Secondly, the z-score of summary statistics contain more statistical information than a sole P value, which might increase the accuracy of the study.

In studies which use real life data, iPheGWAS demonstrated its ability to assemble traits according to their genetic similarity, which, as expected, coincided with their biological domain. Study 1 demonstrated that, even in the presence of negative genetic correlations iPheGWAS performed well. For example, BMI and education was clustered together. Evidence from previous research shows an inverse relationship between education and obesity, albeit weak (27)(28). In study 2, it showed that iPheGWAS was able to cluster traits by genetic similarity instead of by data origination. For example, from Study 2 it was observed that the psychiatric traits and rheumatoid arthritis were located at nearest proximity in the dendrogram indicating these traits were highly correlated. Previous literature shows elevated incidence and prevalence of psychiatric disorders in individuals with rheumatoid arthritis (29)(30). And our finding replicated the finding from LDSC.

## Conclusions

In conclusion, our study has shown that a novel heuristic approach based on the earlier PheGWAS package with an additional hierarchical clustering function is a robust and computationally efficient method for providing information and visualising multiple traits, incorporating information on their genetic correlations.

## Methods

In this study, we applied hierarchical clustering to GWAS summary statistics to identify traits with similar genetic architecture and compared the results to those obtained through LDSC. Firstly, we simulated multiple phenotype data based on varying number of causal SNPs shared between traits, trait heritability and genetic correlation between traits, and used this data to identify the optimal setting of hierarchical clustering for identified genetically similar traits. This was then applied to two real-world datasets to test its performance: (1) 14 traits from the UK Biobank study that were selected to estimate its performance in the presence of positive and negative genetic correlation; and (2) summary statistics from GWAS of 14 phenotypes from various consortiums that selected to estimate its performance when studies included had varying populations, samples sizes, and genetic architecture. The overall study design is shown in figure 1.

### Simulation data

The synthetic data for 10000 individuals with genotypes for 11 million markers which is publicly available in the simulation software portal (https://github.com/precimed/simu) was used for this simulation study. These genotypes were made using 503 European samples from 1000 Genomes Phase 3 (17) data and was produced by running hapgen2 (18).

Two sets of simulation studies were carried out. In set 1, traits were considered as polygenic; 500 SNPs were set as causal SNPs and chosen to be shared across the simulated traits with varying effect sizes. This number was is based on current evidence of polygencity of complex traits, for example for type 2 diabetes there are around 301 to 403 confirmed causal SNPs (19,20). In each round of simulation, 4 traits (namely a, b, c, d) with pre-defined heritability (h^2^) and genetic correlations (rg) were simulated 10 times, under five different proposed genetic architectures (named Arc 1-Arc 5) which varied the h^2^ and genetic correlations between the traits (a, b) and (c, d). The rg between the traits (a,c), (b, d), (a, d), (b, c) was zero throughout the study (Table 1).

In set 2, all settings remained identical except that the number of causal SNPs was set as 60. This was aimed at testing the performance of hierarchical clustering when the traits were less polygenic, as the LDSC has less power in this situation (13).

The simulation was carried out with the software SIMU (21). Further details have been included in the supplementary methods.

### Experimental Setup for Simulation Study

Hierarchical clustering comprises three major stages: (1) feature selection (selecting the elements/features that will be used to carry out the cluster analysis), (2) calculation of distance matrix, and (3) linkage calculations - each stage offering a number of alternative methods of computation.

#### (1) Stage 1 - Feature Selection

In this step, three feature selection methods were tested. For the first two (pairwise and non-pairwise) methods, 32984 LD blocks based on recombination hot spots from Ongen et al (22) were used as feature selection elements. In the third method, the skeleton SNPs defined by LDSC (greater number of features) were used as a fixed set of the feature SNPs.

##### i. Non-pairwise method

For each of the LD Blocks with *p* traits and *q* SNPs, the original matrix was defined as:

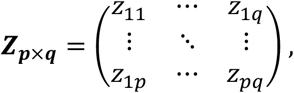

where *z_ij_* represents the z-score for the *i^th^* trait and the *j^th^* SNP, *i* = 1,…, *p* and *j* = 1,…,*q*. The feature selection procedure would select the SNP with the highest absolute z-score corresponding to each trait as the feature SNP, generating *p* feature SNPs. For these *p* feature SNPs, their z-scores corresponding to all *p* traits were then used to construct a *p* × *p* feature matrix ***Y***, defined as below:

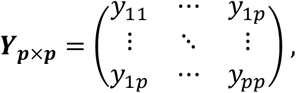

where 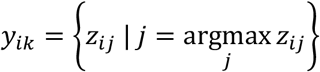 and *k* represents the locations of feature SNPs (one for each trait), *k* = 1,…,*p*

For simplicity, LD Block1 for traits a, b, c, d was selected for illustrating the non-pairwised method in Supplementary figure 1. This process was repeated in all remaining LD blocks. The matrix obtained by combining results of all LD blocks was used to calculate the distance matrix in stage 2.

##### ii. Pairwise method

In this method, separate feature matrices were constructed by considering the traits pairwise instead of considering all traits together as in the non-pairwise method. From an exhaustive combination of the pairs of traits, the non-pairwise approach was applied to these two traits and a 2 × (32984 × 2) z-score matrix was derived for further distance matrix construction in stage 2. This step was repeated 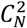 times, where N represents the number of traits. For simplicity, LD Block1 for traits a, b, c, d was selected for illustrating the pairwised method in Supplementary figure 2.

##### iii. Skeleton SNPs

In this method, the 1280026 HapMap3 common SNPs (derived from website https://www.sanger.ac.uk/resources/downloads/human/hapmap3.html), called ‘Skeleton SNPs’ were used for defining the features. The z-score corresponding to each of these skeleton SNPs was used to construct the feature matrix that in turn was used to calculate the distance matrix in stage 2.

#### (2) Stage 2 -Calculation of distance matrix

From the feature matrices generated in the first stage, the distance matrix was calculated using the Pearson and Spearman correlation methods. The absolute values of the correlation matrix were used to capture the strength of both negative and positive correlation and obtain the distance matrix.

#### (3) Stage 3 -Linkage calculations

In this stage the Average distance method was used for linkage. This method uses average pairwise distances between components that are to be merged into clusters.

### Evaluation of the simulation method

Each simulation result was considered to be valid if the ground truth was observed in the results, I.e. traits (a, b) and the traits (c, d) clustered together. In situations where (a,b) and (c,d) fail to cluster together, the results were considered to be unsuccessful. For comparison, LDSC was used to generate the genetic correlation coefficients for different pairs of traits and hierarchical clustering was applied to the correlation matrix. The method with the highest successful rate was chosen.

### Real world data study 1-Traits selected from UKBB

This study was designed to estimate the sensitivity of iPheGWAS to ordering and clustering traits given a known large number of positive and negative correlations within a population. Within UKBB, 14 traits from various domains were tested, including ‘self-harm’, ‘neuroticism score’, ‘mood swing’ from the mental health domain; ‘pain in throat or chest’, ‘atherosclerosis’, ‘no heart problem’, ‘cholesterol lowering’ and ‘ECG load’ from the cardiology domain; traits from anthropomorphic domain (‘stand height’, ‘trunk’, ‘mass’) and some food related traits (‘portion size’, ‘food weight’ and ‘potassium’).

### Real world data study 2-Traits selected from various consortiums

In study 2, traits were selected from different consortiums to investigate the performance of iPheGWAS when traits were chosen from different study populations. Summary statistics from GWAS of 14 phenotypes, including five psychiatric traits (‘attention deficit hyperactivity’, ‘major depressive’, ‘autism’, ‘bipolar disorder’, ‘schizophrenia’) and four autoimmune diseases (‘rheumatoid arthritis’, ‘ulcerative colitis’, ‘inflammatory bowel disease’,’Crohn’s disease’), four anthropomorphic traits (‘waist hip ratio’, ‘BMI’, ‘height’, ‘birth weight’) and educational attainment were used.

### Real world data validation

iPheGWAS was applied to the two sets of real-world datasets. Optimized linear ordering (OLO) (23) was then applied to the leaves in the clusters of the dendrogram to minimize the distance between successive clusters. As a comparison, LDSC was used to generate the genetic correlation coefficients for different pairs of traits and hierarchical clustering was applied to the correlation matrix to generate a dendrogram.

For iPheGWAS methods, internal validation was performed by comparing the correlation between the cophenetic distances and the original distance data (cophenetic correlation coefficient), in order to indicate how well the dendrograms represent the real data. A more precise solution is believed to be obtained as the correlation coefficient moves closer to 1.

For external validation, Tanglegram was used to visually inspect the structure of dendrograms derived from iPheGWAS and LDSC, and the entanglement coefficient was used to assess the quality of the alignment of two trees. Entanglement values ranged between 0 and 1 with a lower entanglement coefficient corresponding to a good alignment. Tree based external validation was also performed and “Baker” and “Cophenetic” correlation matrix were computed between trees. Values near 0 indicated that two trees are not statistically similar.

Rand Index (RI) and Adjusted Rand Index (ARI) were used to measure similarity between iPheGWAS clustering and LDSC clustering. The higher the ARI, there is higher agreement between two clusters with a maximum value of 1.

### Implementation

The hierarchical clustering was performed using the R *stats* package (version 4.0.1). The *factoextra* package in R was used to plot the dendrogram.

The code for this heuristic approach has been scripted in R version 4.0.1 and it has been incorporated into the PheGWAS package, named iPheGWAS. An external matrix module was also added which makes use of the genetic correlation matrix generated by other methods for inspecting the genetic correlation values.

## Availability of data and materials

The iPheGWAS software and code are freely available in the GitHub repository (https://github.com/georgeg0/iphegwas). The details and the guidance of the software package can be found in https://georgeg0.github.io/iphegwas/index.html.

The datasets supporting the conclusions of this article are available as follow:

Psychiatric Genomics Consortium, https://www.med.unc.edu/pgc/results-and-downloads/downloads;

SSGAC (educational attainment), https://www.thessgac.org/data;

GIANT Consortium, https://portals.broadinstitute.org/collaboration/giant;

Early Growth Genetics Consortium, http://egg-consortium.org/birth-weight-2016.html;

Rheumatoid arthritis, http://plaza.umin.ac.jp/~yokada/datasource/software.htm;

International Inflammatory Bowel Disease Genetics Consortium, https://www.ibdgenetics.org/.

## Supplementary Data

Supplemental data include 1 supplementary method section, 3 supplementary results section and 6 figures.

## Acknowledgement

We would like to thank the Health Informatics Centre (HIC), the principal investigators and our colleagues in the INSPIRED project.

## Funding

This work was supported by the National Institute for Health Research using Official Development Assistance (ODA) funding [INSPIRED 16/136/102].

## Conflict of Interest

None declared.

## Contributions

G.G., Y.H. and A.D. conceived and designed the study. G.G., Y.H., A.B. and H.W. developed the algorithms. G.G., S.G., and A.N. performed the analysis. G.G. implemented the software package. G.G., and Y.H. prepared the original draft. H.W., A.B., C.P. and A.D. provided critical feedback on the design application and implementation of the tool. S.G., A.N., J.H., R.V., V.M, and A.D. helped with the manuscript revising. The authors read and approved the final manuscript.

## Notes

### Competing Interest Statement

The authors have declared no competing interest.

