## Supplementary File for "iPheGWAS: an intelligent computational framework to integrate and visualise genome-phenome wide association results"

**Supplementary Methods:**

**Steps involved in Simulation**

The following steps were undertaken to conduct the simulation:

i) Genetic covariance matrix that was generated based on the pre -defined heritability and genetic correlations.

ii) A list of causal SNPs were picked randomly

iii) For the selected causal SNPs, simple additive effects were drawn from the multivariate normal distribution based on the genetic covariance matrix (obtained from step i).

iv) The causal SNPs along with the true effects in combination with the synthetic genotype were used in SIMU to generate synthetic phenotypes a, b, c, d. Here I have used the same set of randomly selected causal SNPs for generating the phenotypes to set a scenario of polygenic overlap.

v) A GWAS was performed using synthetic phenotypic data and genotype data to obtain the summary statistics.

**Supplementary Results 1**

**Real world data validation: Study 1- Traits selected from UKBB (iPheGWAS)**

it was observed that the leaves corresponding to ‘trunk’ and ‘mass’ have the shortest height indicating they were the closest and got clustered first (clade 1). Following closely, ‘neuroticism score’ and ‘mood swing' clustered (clade 2) and the leaves 'food weight' and 'potassium' clustered (clade 3). Although the intra-element distance in these two clusters were small, suggesting higher genetic correlation within domains, it is to be noted that the inter cluster distances between them is considerably large, placing them at two extreme ends of the dendrogram. Next, 'stand height' was found to cluster with 'trunk' and 'mass' (clade 4) showing that it was closer to this cluster than with any other element in the tree. Next, 'cholesterol lowering' and 'no heart problem' formed a cluster (clade 5). This formed one of the two branches of the clade 7, of which the other branch was formed by the leaves 'pain throat' and 'atherosclerosis' (clade 6). Clade 2 was joined by a simplicifolius 'self harm' (clade 8). Further, 'portion size' clustered with cluster 'trunk', 'mass' and 'stand height' (clade 9). 'ECG load' clustered with the other cardiology traits (clade 10). Clade 8 and clade 10 joined to form a cluster (clade 12) which finally joined clade 11.

**Supplementary Results 2**

**Real world data validation: Study 1- Traits selected from UKBB (LDSC)**

The dendrogram for study 1, LDSC method is presented in Figure 5. For external tree based validation with the dendrogram from genetic correlations calculated by LDSC, 'neuroticism score' and 'mood swings' clustered first similar to our method and 'self harm' was found to join this cluster as a simplicifolius (clade 8)- but in LDSC method 'self harm' was found to be located closer to 'neuroticism score'. 'potassium' and 'food weight' clustered secondly, the order of the leaves remaining same as obtained from the heuristic method. 'mass' and 'trunk' clustered thirdly and the leaf 'portion size' was found to be the closest to these cluster (clade 5) followed by 'stand height' (clade 9) which joined as a simplicifolius. 'ECG load' and 'pain in throat/chest' were found to be more similar unlike in our method and these formed a cluster (clade 4). ‘No heart problem’ and ‘cholesterol lowering’ were found to be closer to each other forming a cluster and ‘atherosclerosis’ joined as a singleton to this cluster (clade 6). The cluster formed by clade 2 was found to be closer to the cluster formed by clade 9 and these two clusters were joined by clade 12 to form a bigger cluster. On the other hand, the clusters formed from clade 8 and 10 were found to be more similar and these joined to form a bigger cluster(clade11). Finally, clade 11 and 12 joined to form a single big cluster.

**Supplementary Results 3**

**Real world data validation: Study 2- Traits selected from various consortiums (iPheGWAS)**

In study 2, it is observed that clade 1 has five leaves representing the psychiatry traits. The lowest height indicates that these traits of clade 1 are more similar to each other as compared to any other traits in the tree and were clustered first. Next, we move to clade 2, which has two branches- the first branch with the five psychiatry leaves and second branch with a single leaf representing ‘Rheumatoid Arthritis’ (cluster 1). From the height of the leaves, it is observed that the leaves ‘Ulcerative Colitis’ and ‘IBD’ (Clade 3) were clustered next. These leaves are joined in clade 4 to ‘Crohns Disease’. Following this it is found that ‘waist hip ratio’ and bmi cluster together (clade 5). Following this, birthweight and height cluster (clade 6). Education is found to be more similar to ‘waist hip ratio’ and ‘BMI’ and joined as a simplicifolius to this cluster by clade 7. After this, anthropomorphic traits converged to form clade 8. Further, clade 8 joined clade 4 to form a bigger cluster (Cluster 2) and finally, Cluster 1 joined the Cluster 2 through clade 9.

**Supplementary Figures:**

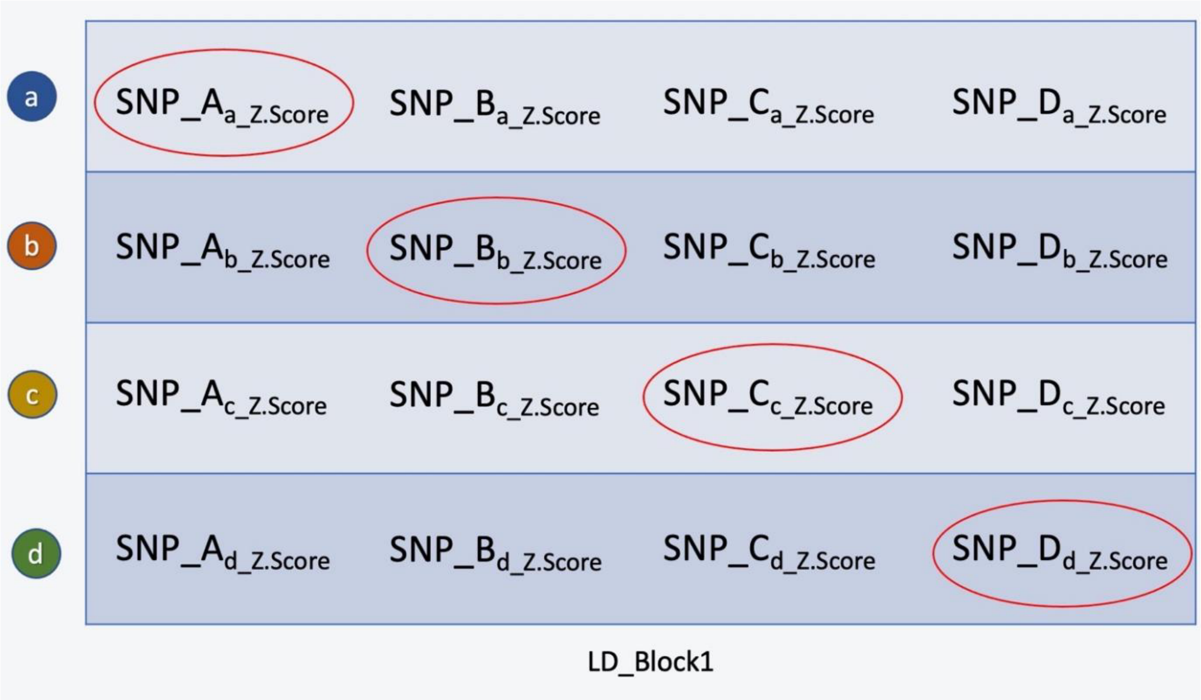


**Supplementary figure 1.** Illustration explaining the non-pairwise method: for each trait, the feature SNPs from this block based on the highest absolute z-score corresponding to each of the traits (circled in red). Corresponding to trait 'a', 'SNP_A' was found to have the highest absolute z-score in LD_Block1. This SNP was selected as one of the feature SNPs. The z-score of 'SNP_A' was recorded for all the other traits(b,c,d). This is repeated for all the traits and a N by N (number of traits) feature matrix is constructed for LD_Block1. This process occurs in all LD blocks simultaneously and a feature matrix combining results from all LD blocks is obtained. Therefore, for each trait, total number of feature SNPs obtained is 32984 × N (number of LD blocks), and the matrix for hierarchichal clustering is N traits by 32984 × N feature SNPs. (In some LD blocks the SNPs corresponding to the highest z-score may coincide for two or more traits)

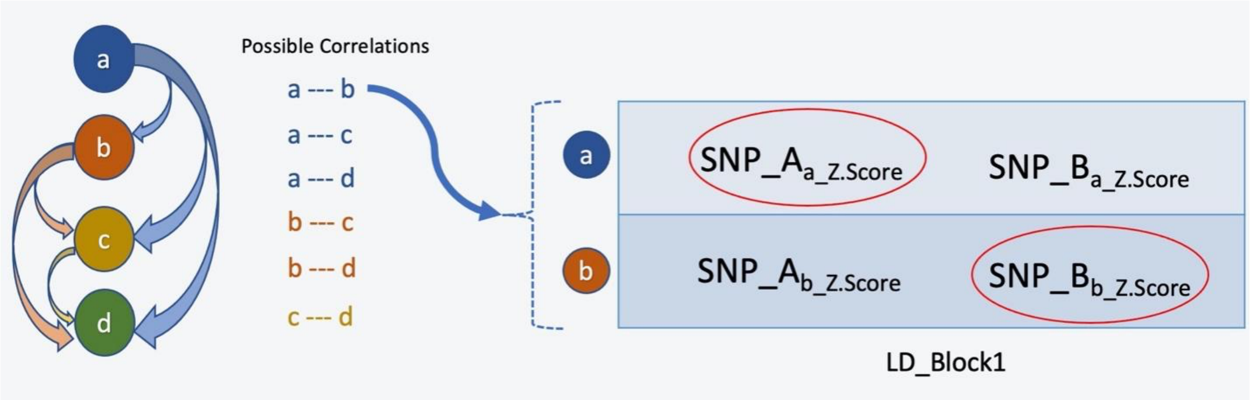


**Supplementary figure 2.** Illustration explaining the pairwise method: each time, a pair of traits were tested. For example, trait a and b were test for the first round. Corresponding to trait ‘a’, ‘SNP_A’ was found to have the highest absolute z-score in LD_Block1. This SNP was selected as one of the feature SNPs. The z-score of ‘SNP_A’ was recorded for the paired trait b. Same for trait b a 2 by 2 feature matrix is constructed for LD_Block1. This process occurs in all LD blocks simultaneously and a feature matrix combining results from all LD blocks is obtained. Therefore, for each trait, total number of feature SNPs obtained is 32984 × 2 (number of LD blocks), and the matrix for hierarchichal clustering is 2 traits by 32984 × 2 feature SNPs. (In some LD blocks the SNPs corresponding to the highest z-score may coincide for two or more traits). After tested for the first pair of trait, the other possible pairs of combinations were tested in the following rounds and the overall rounds of tests are $C_N^2$ times, where N represents the number of traits.

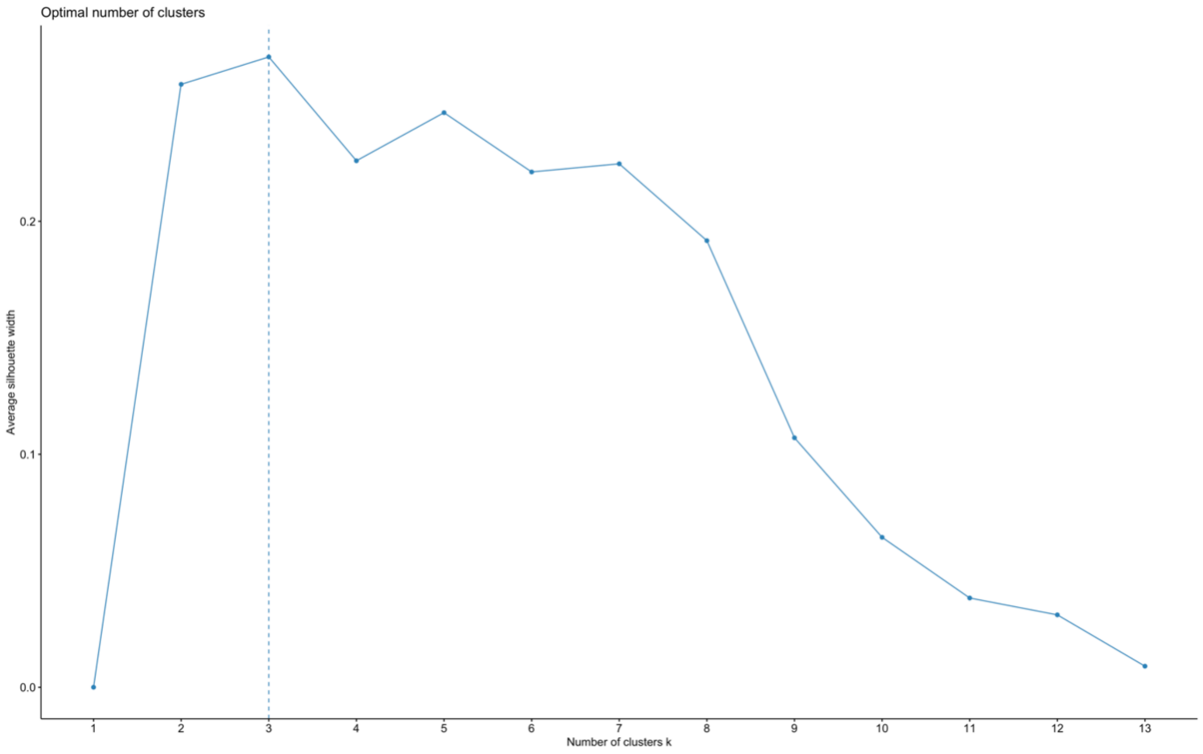


**Supplementary figure 3.** Silhouette plot representing the optimal number of clusters for study 1 using iPheGWAS

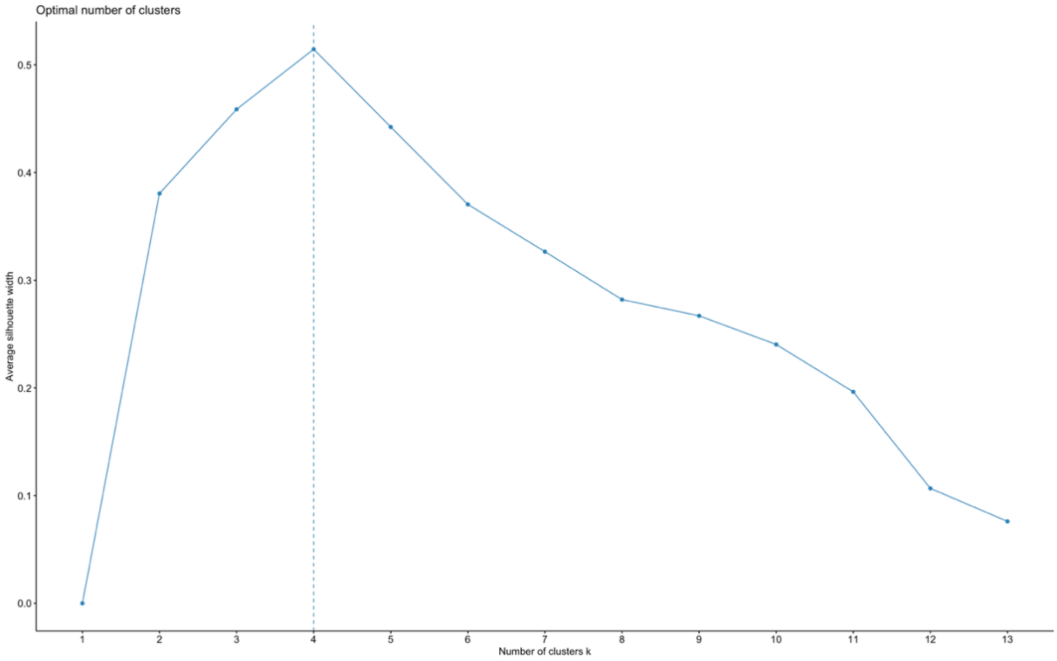


**Supplementary figure 4.** Silhouette plot representing the optimal number of clusters for study 2 using LDSC method.

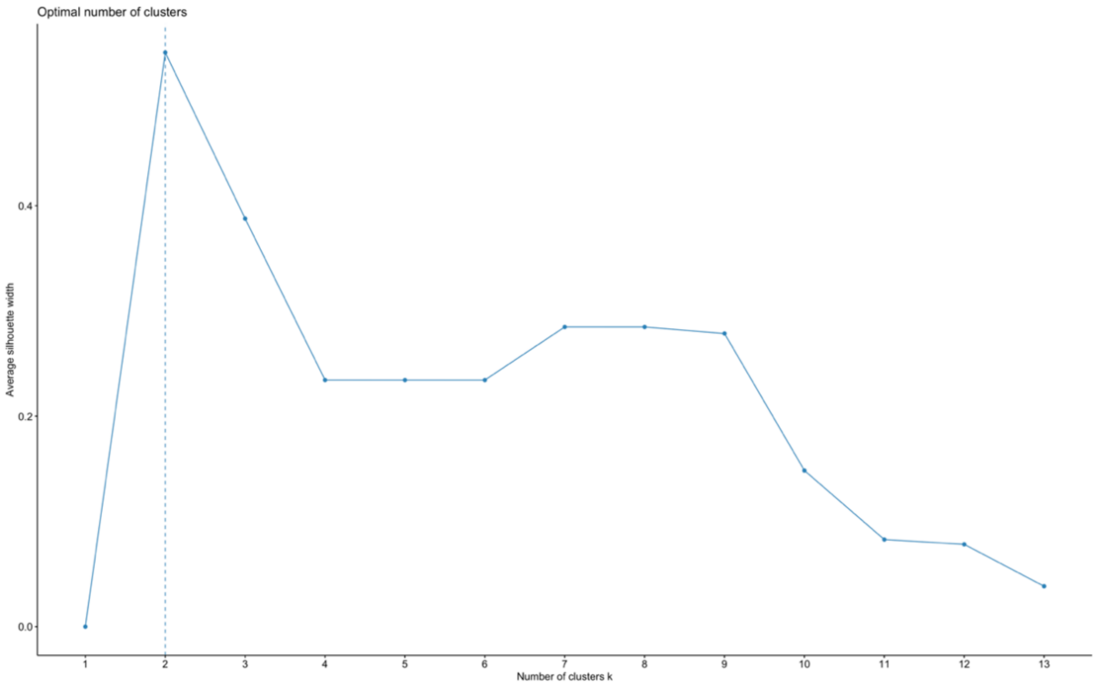


**Supplementary figure 5.** Silhouette plot representing the optimal number of clusters for study 2 using our method.

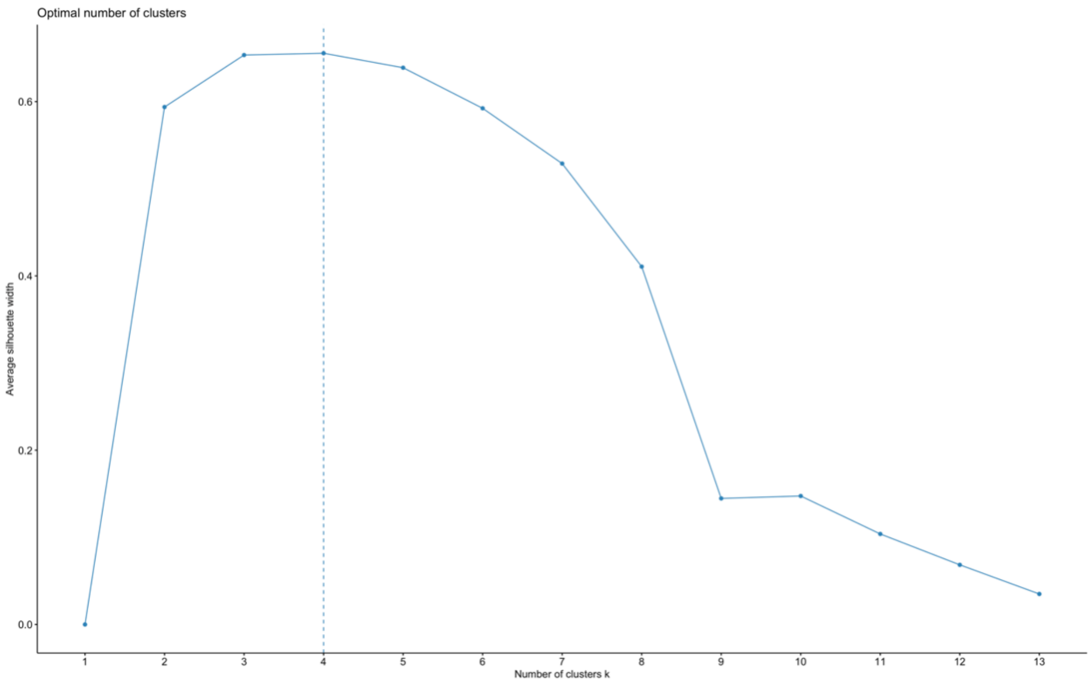


**Supplementary figure 6.** Silhouette plot representing the optimal number of clusters for study 2 using LDSC method.
